# Protoporphyrin IX is a dual inhibitor of p53/MDM2 and p53/MDM4 interactions and induces apoptosis in B-cell chronic lymphocytic leukaemia cells

**DOI:** 10.1101/548875

**Authors:** Liren Jiang, Natasha Malik, Pilar Acedo, Joanna Zawacka-Pankau

## Abstract

p53 is a tumor suppressor, which belongs to the p53 family of proteins. The family consists of p53, p63 and p73 proteins, which share similar structure and function. Activation of wild-type p53 or TAp73 in tumors leads to tumor regression, and small molecules restoring the p53 pathway are in clinical development.

Protoporphyrin IX (PpIX), a metabolite of aminolevulinic acid, is a clinically approved drug applied in photodynamic diagnosis and therapy. PpIX induces p53- and TAp73-dependent apoptosis and inhibits TAp73/MDM2 and TAp73/MDM4 interactions. Here we demonstrate that PpIX is a dual inhibitor of p53/MDM2 and p53/MDM4 interactions and activates apoptosis in B-cell chronic lymphocytic leukaemia cells without illumination and without affecting normal cells. PpIX stabilizes p53 and TAp73 proteins, induces p53-downstream apoptotic targets and provokes cancer cell death at doses non-toxic to normal cells.

Our findings open up new opportunities for repurposing PpIX for treating lymphoblastic leukaemias with *wtTP53*.

## Introduction

B-cell chronic lymphocytic leukaemia (CLL) is one of the most common forms of blood cancers^1,2^. The incidence of CLL in the western world is 4.2/100 000 per year. In patients over the age of 80 years, the incidence increases to greater than 30/100 000 per year. CLL develops mostly in patients above the age of 72 years which is linked to poor prognosis^3^.

In Sweden, and worldwide, male show higher prevalence of lymphoid and haematological tissue cancers than women and the leukaemia incidence in both genders increases above the age of 55^4^.

Common chromosomal aberrations in CLL include deletions of 13q14, trisomy 12, 11q23 and 17p13 deletions or mutations^5^. Deletions of 17p13 and 11q23 affect p53 pathway and are linked with poor prognosis and faster disease progression^6^. Recent studies point to the positive outcome of the Bcl-2 inhibitor venetoclax in handling 17p-deleted relapsed/refractory CLL^7^. The clinical trial for the treatment-näive CLL elderly patients show promise for Ibrutinib, a Bruton’s tyrosine kinase (BTK) inhibitor^8^. However, the development of resistance to targeted therapies poses significant therapeutic constraints.

Introduction of new compounds into clinical practice is both, time constraining and a financial endeavour, which more often than not is subject to failure. Drug repurposing brings a selective advantage to the field of drug discovery as it is easier and more cost-effective to authorize the approved drug for a new indication^9^.

Protoporphyrin IX (PpIX) is a natural precursor of heme and a metabolite of aminolevulinic acid, a photoactivable drug clinically used in photodynamic therapy and diagnosis^10^. In drug repurposing approach PpIX was identified as an activator of p53 and TAp73α tumor suppressors. Recent work demonstrates that PpIX inhibits TAp73/MDM2 and TAp73/MDM4 interactions, which leads to stabilisation of TAp73 protein and induction of TAp73-dependent apoptosis in cancers with *TP53* gene mutations^11,12^.

The tumor suppressor p53 is inactivated in the majority of tumors by mutations occurring in the *TP53* gene^13^. In cancers retaining intact *TP53* gene, p53 protein is targeted for degradation by the deregulated E3 ubiquitin ligase MDM2. In addition, MDM2 homolog, MDM4, protein binds p53 and inhibits its transcription activity^14 15 16^.

Activation of wild-type (wt) p53 is a promising therapeutic strategy, and the compounds inhibiting oncogenic MDM2 or modulating p53 post-translational modifications are currently in the clinical development^17^. However, due to systemic toxicity, highly selective inhibitors of p53/MDM2 interactions including analogues of nutlin, MI or RG compounds, have not been approved yet^18 19^. Even though the advancement in the field, these compounds cannot inhibit MDM4 protein and are thus inefficient in targeting tumors that overexpress MDM4 oncogene such as cutaneous melanomas^20^.

p73 is a tumor suppressor and induces apoptosis and tumor regression in a p53-independent manner^21–23^. *TP73* gene is rarely mutated in cancers and p73 protein is often inactivated by binding to oncogenic partners including MDM2, MDM4, ΔNp73 or mutant p53^24^. Strategies aiming at targeted activation of p73 in cancer are, however, at a very early stage of development.

Here, we applied a fluorescent two-hybrid assay and a yeast-based reporter assay and showed that PpIX inhibits p53/MDM2 and p53/MDM4 interactions. Next, analysis in cancer cells revealed that PpIX induces the p53-dependent apoptosis in B-CLL cells. We demonstrate that PpIX triggers accumulation of p53 and TAp73 and activates cell death at doses not affecting healthy peripheral blood mononuclear cells (PBMCs).

## Materials and methods

### Reagents and cell lines

Protoporphyrin IX and nutlin were purchased from Sigma-Aldrich (Munich, Germany) and re-constituted in 100% DMSO (Sigma-Aldrich, Munich, Germany) to 2 mg/ml or 10 mM respectively. PpIX was stored in amber eppendorf tubes at room temperature and nutlin was aliquoted and stored at −20°C. RITA was purchased from Calbiochem (Solna, Sweden) reconstituted in 100% DMSO to 0.1 M, aliquoted and stored at −20°C.

Cisplatin (Sigma-Aldrich, Munich, Germany) was prepared in 0.9% NaCl solution to 1mM, protected from light and stored at −20°C. MG132 was from Sigma-Aldrich (Munich, Germany) reconstituted in 100% DMSO to 10 mM and stored at −20 °C. IgG and protein A agarose beads were from Santa Cruz Biotechnology (Solna, Sweden), protease inhibitors were prepared from tablets oComplete® Roche to 100x concentration (Sigma-Aldrich, Munich, Germany), MTT was from Sigma-Aldrich (Munich, Germany).

Rabbit polyclonal anti-MDMX was from Imgenex (Cambridge, UK), rabbit polyclonal anti-TAp73 (A300-126A) (Bethyl Laboratories, TX, USA), anti-PUMA (ABC158; Merck, MA, USA), anti-BAX (N-20; Santa Cruz Biotechnology, Germany), anti-Bid (FL-195; Santa Cruz Biotechnology, TX, USA), anti-PARP (F-2; Santa Cruz Biotechnology), anti-β-actin (A2228; Sigma-Aldrich), normal mouse IgG (sc-2025) were from Santa Cruz Biotechnology. Antimouse HRP and anti-rabbit HRP secondary antibodies were from (Jackson ImmunoResearch Inc.) Reverse transcription iScript cDNA synthesis kit and SSo Advanced Universal SYBR Green kit were from Bio-Rad (Solna, Sweden)^25^.

### Cell lines

EHEB (wt-p53) chronic B cell leukaemia cells were kindly provided by Dr Anders Österborg, Karolinska Institutet (source ATCC). HL-60 (p53-null) acute promyelocytic leukemia cell lines were provided by Dr Lars-Gunnar Larsson, Karolinska Institutet (source ATCC). PBMCs were provided by Dr Noemi Nagy, Karolinska Institutet and separated as described previously^26^. HCT 116 cells were a kind gift from Dr Bert Vogelstein^27^.

Leukemic cells and PBMCs were cultured in RPMI-1640 (Roswell Park Memorial Institute) medium (Sigma-Aldrich, Munich, Germany) and HCT 116 cells in DMEM medium with 10% foetal calf serum (Sigma-Aldrich) and penicillin/streptomycin (10 units/ml) (Sigma-Aldrich) at 37 °C in a humidified 5% CO_2_/95% air atmosphere.

### Cell Viability Assay

The viability of EHEB, HL60 and PBMCs after 72-hour treatment with PpIX was assessed by the 3-(4,5-dimethylthiazol-2-yl)-2,5-diphenyltetrazolium bromide (MTT) assay according to manufacturer’s protocol. Briefly, 5mg/ml MTT solution was prepared in PBS buffer and filter-sterilized. Cells were washed once with RPMI-1640 medium and 1*10^5^ cells/ml were transferred to eppendorf tubes and treated with 0.5%DMSO or the investigated compounds. Next, cells were seeded onto 96 well plates at the density of 1*10^4^ cells/well and incubated for 72h at 37 °C. After this time, MTT reagent was added to each well to a final concentration of 10% and the plates were incubated for 3h at 37 °C in a humidified 5% CO_2_/95% air atmosphere. The supernatant was removed and 200 μl DMSO/well was added. The plates were incubated at 37°C for 30 min and the absorbance of the formazan was measured at 560 nm in a Perkin-Elmer (Waltham, MA, USA) microplate reader.

Untreated EHEB cells, RPMI 1640 medium and RPMI 1640 medium with 20μg/ml PpIX were used as background controls. The experiments were performed in triplicates and in at least three independent repeats.

### Immunoprecipitation and western blot

Immunoprecipitation was performed using a modified protocol described previously^28^. Briefly, HCT 116 cells were seeded in 10 cm dish at 4*10^6^ cells, allowed to adhere overnight and treated with the compounds for 8h followed by 3h with 20 μM MG132. Cells were washed 2x with ice-cold PBS and solubilized in IP buffer: 25 mM Tris-HCl, pH 8.0, 150 mM NaCl, and 0.5% Nonidet P-40 supplemented with protease inhibitors and lyzed on ice for 30 min and 1 mg of total protein in IP buffer was added to 30 μl mouse protein A agarose beads and 1 μg mouse anti-p53 DO-1 antibody or 1 μg normal mouse IgG and immunoprecipitated for 16h at 4°C. The beads were washed three times with buffer 1 (50 mM Tris-HCl, pH 7.5, 5 mM EDTA, 500 mM NaCl, and 0.5% Nonidet P-40) and one time with buffer 2 (50 mM Tris-HCl pH 7.5, 5 mM EDTA, 150 mM NaCl). The beads were resuspended in 15 μl of lysis buffer and 5 μl of 5 × Laemmli buffer and boiled prior to western blot analysis.

For western blot, total proteins were transferred to HyBond membrane (GE healthcare), blocked with 5% milk in PBS for 20min and incubated with the relevant antibodies. After washing in PBS membranes were incubated with secondary antibodies (1:3000 in 5% milk) for 2h at room temperature. The protein signals were detected using Super Signal West Dura Extended Duration Substrate (Bio-Rad, Solna, Sweden) and ChemiDoc (Bio-Rad).

PBMCs were cultured in RPMI 1640 supplemented with 10% foetal calf serum and penicillin/streptomycin (10 units/ml) without supplementation with growth factors at 37 °C for three days before treatment with compounds.

### Quantitative PCR

Quantitative PCR was performed as described previously^25,29^. Briefly, cells were treated with 2.5 μg/ml PpIX for 8h. qPCR was performed using following primers pairs:

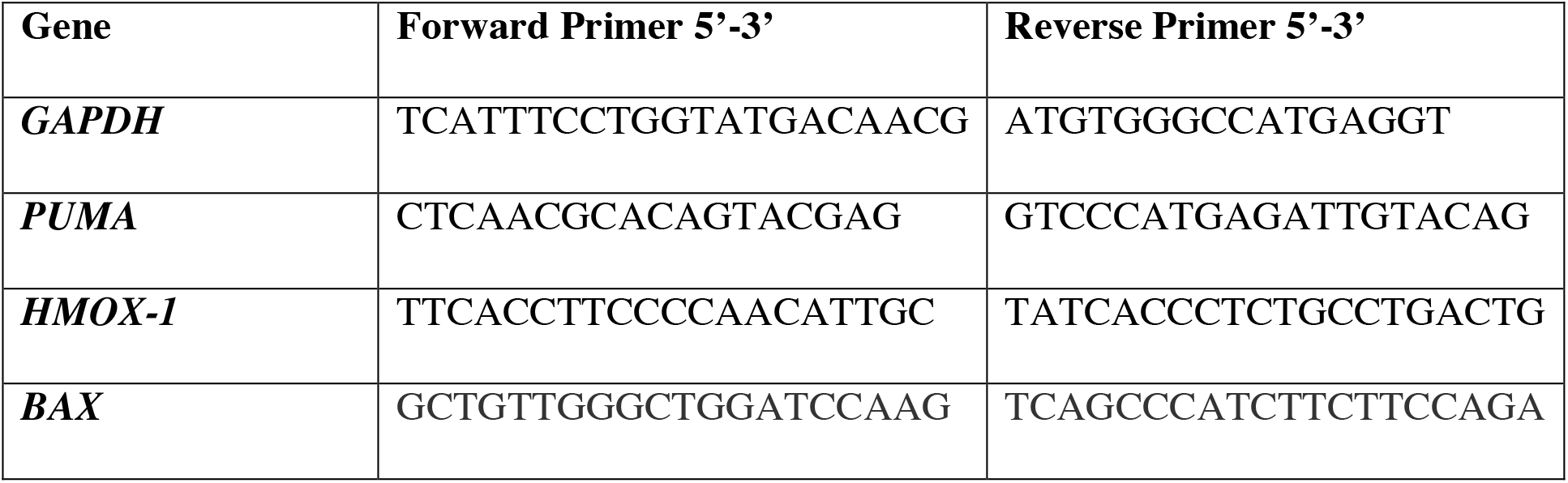

### Fluorescence Activated Cell Sorting (FACS)

Cells were cultured in 6-well plates with 8*10^5^ (K562 and HL60) and 1*10^6^ PBMC cells and 2 ml media/well and treated with the compounds. Propidium iodide (PI) and FITC-Annexin V (both from BD Biosciences, CA, USA) staining was performed according to the manufacturer’s protocols. For PBMCs, cells were washed and fixed with 500 μl ice-cold 70% ethanol and stored at 4°C. Next, cells were washed and stained in 300 μl PI solution, stained for 30 min, spanned down and suspended in 200 μl PBS. FACS was carried out using the CELLQuest software (CELLQuest, NJ, USA) as described previously^30^.

#### Yeast-based reporter assay

The yeast-based functional assay was conducted as previously described^31^. Briefly, the p53-dependent yeast reporter strain yLFM-PUMA containing the luciferase cDNA cloned at the *ADE2* locus and expressed under the control of PUMA promoter^32^ was transfected with pTSG-p53^33^, pRB-MDM2 (generously provided by Dr. R. Brachmann, Univ. of California, Irvine, CA, USA), or pTSG-p53 S33/37 mutant and selected on double drop-out media for TRP1 and HIS3. Luciferase activity was measured 16h after the shift to galactose-containing media^32^ and the addition of 2 and 10 μg/ml PpIX or 10 or 50 μM nutlin (Alexis Biochemicals, Sant Diego, CA, USA), or DMSO. Presented are average relative light units and the standard errors obtained from three independent experiments each containing five biological repeats.

#### F2H®-analysis

The assay was developed and performed as described previously^34,35^. Briefly, F2H®-analysis (ChromoTek GmbH, Planneg-Martinsried, Germany) was carried out to assess PpIX ability to disrupt p53/MDM2 and p53/MDM4 interactions in U2OS cells, when MDM2 or MDM4 was tethered in the nucleus. U2OS cells were co-transfected with LacI-GFP-MDM2 or MDM4 and RFP-p53 for 8h and then incubated with 1 or 10 μM of PpIX or nutlin for 16h. Control interaction values in each independent experiment are normalized to 100%. Averaged interaction values for the treated cells were plotted for p53/MDM2 and p53/MDM4 interactions on the graph. Data is presented as mean ± s.e.m., n = 6, PpIX - p < 0.01, nutlin - p < 0.001, Student’s *t*-test.

## Results

### PpIX ablates p53/MDM2 and p53/MDM4 interaction

It has been previously shown that PpIX inhibits p53/MDM2 interactions, induces p53-dependent reporter and apoptosis in human cancer cells expressing wild-type p53^36,37^. Next, PpIX was described to inhibit TAp73/MDM2 and TAp73/MDM4 interactions and to activate the TAp73-dependent apoptosis in cancer cells harbouring mutant *TP53* gene^29 12^. The mechanism of inhibition of protein-protein interactions (PPIs) is via binding of PpIX to the N-terminus of TAp73^37^. Thus, here, we strived to investigate if PpIX, which unlike nutlin, binds to the N-terminus of p53^37^ and not to MDM2, is also capable of inhibiting the interaction between p53 and MDM4. We engaged the Fluorescent Two-Hybrid (F2H®) analysis performed by ChromoTek GmbH in which the GFP-labelled MDM2 or MDM4 proteins (LacI-GFP-MDM2 or MDM4) were tethered at the nucleus of U2OS cells and examined for the localisation of the exogenously expressed RFP-labelled p53 protein before and after drug treatment for 16h. The data were analysed using fluorescent imaging as described previously^35^. We observed an inhibitory effect of 10 μM PpIX on both interactions – p53/MDM2 and p53/MDM4. Interaction values dropped from 100% in untreated cells down to 61 ± 8 % for p53/MDM2 (mean ± s.e.m, Student’s t-test, p < 0.01, n = 6) and 79 ± 5 % for p53/MDM4 (p < 0.05, n = 6). For comparison, the reference compound, 10 μM nutlin-3, induced a similar reduction of the p53/MDM2 interaction (down to 60%), along with reductions of the p53/MDM2 interactions at a lower dose of 1 μM (p < 0.001, Student’s t-test, n = 6). Nutlin-3 did not disrupt the p53/MDM4 interaction (**Figure 1A**).

**Figure 1.**
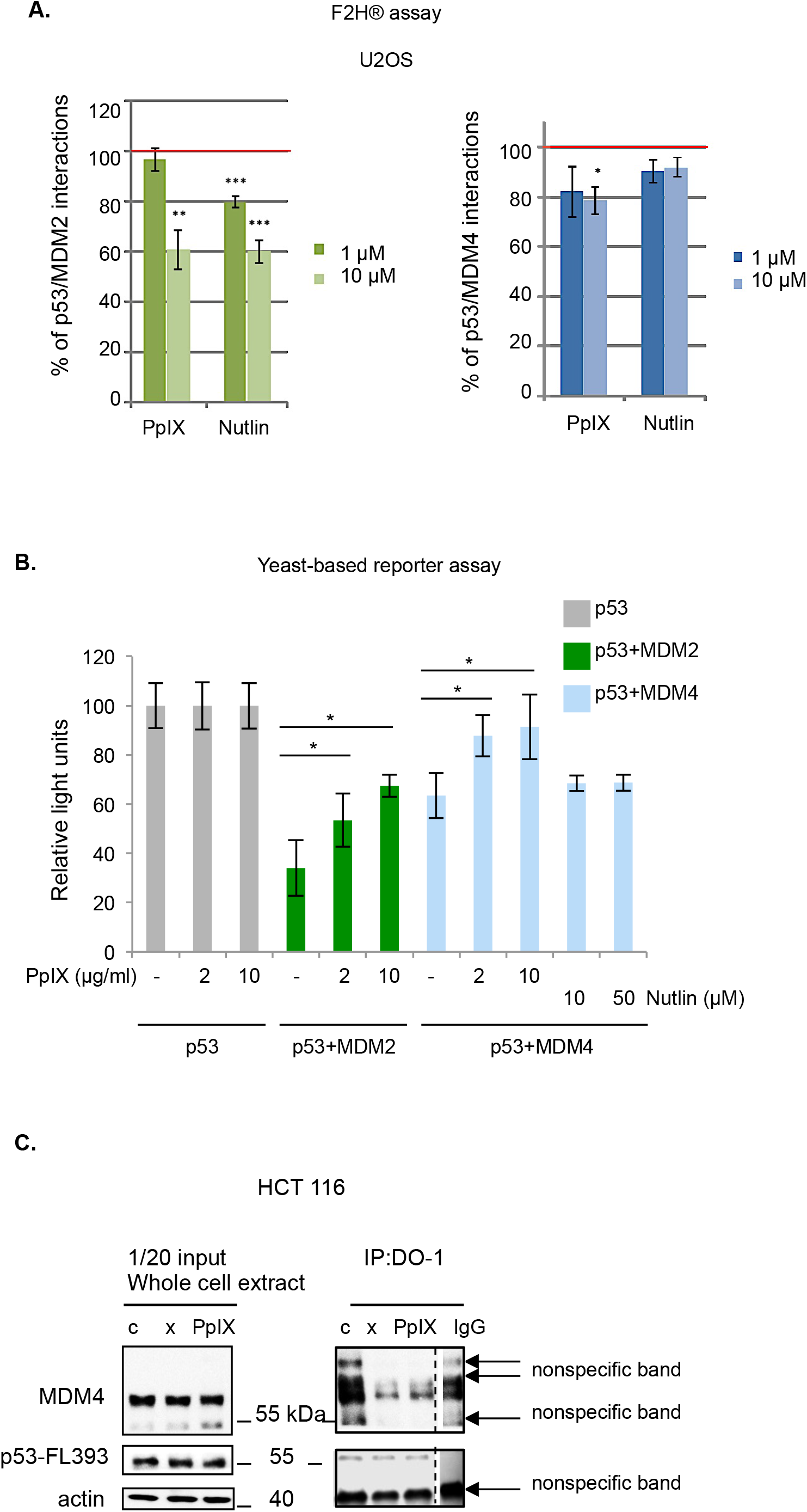
Protoporphyrin IX inhibits p53/MDM2 and p53/MDM4 interactions. A. F2H®-analysis of disruption of interactions of p53/MDM2 and p53/MDM4 by protoporphyrin IX (PpIX) and nutlin in U2OS cells, when MDM2 or MDM4 is tethered in the nucleus. U2OS cells were co-transfected with LacI-GFP-MDM2 or MDM4 and RFP-p53 for 8h and then treated with PpIX or nutlin for 16h. Control interaction values are normalized to 100% and are shown as a red line. Averaged interaction values for the treated cells are plotted for p53/MDM2 interaction (left, green bars) and p53/MDM4 interaction (right, blue bars). Data represented as mean ± s.e.m., n = 5 - 6, *-p < 0.05, ** - p < 0.01, *** - p < 0.001, Student’s t-test. B. Rescue of the p53 transcription activity in the presence of MDM4 by PpIX but not nutlin as assessed in the yeast-based reporter assay. The average light units relative to the transactivation activity of p53 alone and the standard errors of at least five biological repeats are presented. The Student’s *t*-test was performed for statistical analysis with p ≤ 0.05. C. PpIX (1 μg/ml) disrupts p53/MDM4 interactions in HCT 116 colon cancer cells as assessed by immunoprecipitation. The data is a representative of two independent experiments. Cropped line represents the site were the membranes were cut. X-sample not relevant to this study.

Next, to determine that the inhibition of p53/MDM2 and p53/MDM4 complexes by PpIX is due to the direct interaction between the compound and p53 and not due to post-translational modifications in the p53 protein, we employed a yeast-based reporter assay which allows measuring the transcription activity of human p53 using p53-dependent luciferase reporter^11,31^. Since p53 is not degraded by the exogenous human MDM2 or MDM4 in yeast cells, the inhibitory effect of MDM2 in this system can be ascribed to the direct interaction with p53 and the interference with the p53-dependent gene transcription^38^. Cotransfection of MDM2 inhibited the activity of p53 as expressed by the drop from 100 relative light units for p53 itself to 34.02 relative light units in the presence of MDM2 (**Figure 1B**). 2 and 10 μg/ml PpIX rescued the wt-p53-mediated activation of luciferase expression to 53.42 and 67.39 relative light units respectively. The effect on the p53/MDM4 interactions was also significant; PpIX increased the relative light units from 63.48 to 87.79 and 91.42 at 2 and 10 μg/ml respectively. The reference compound, nutlin-3 did not rescue the transcription activity of p53 in the presence of MDM4 indicating that it does not inhibit p53/MDM4 interactions.

To determine that PpIX inhibits p53/MDM4 interactions also in cancer cells, we treated HCT 116 human colon cancer cells with PpIX for 16h. To stabilize MDM4 that is otherwise driven to degradation by MDM2 released from the complex with p53 by PpIX, cells were treated with MG132. Immunoprecipitation with monoclonal anti-p53 DO-1 antibody was performed, and the membrane was blotted with the polyclonal anti-p53 FL-393. The results demonstrate that PpIX readily inhibited the interaction between p53 and MDM4 (**Figure 1C**). We detected unspecific binding of MDM4 to mouse IgG, however, the signal was much weaker than the one obtained for the untreated cells (**Figure 1C**), thus the background binding does not affect the conclusion that PpIX readily inhibits p53/MDM4 interactions in cancer cells.

Thus, F2H® yeast-based reporter assay and pull down demonstrated that the binding of PpIX to p53 results in the inhibition of both p53/MDM2 and p53/MDM4 interactions.

#### PpIX inhibits proliferation and induces apoptosis in B-CLL leukaemia cells

PpIX was shown to induce apoptosis^39^, p53-, p73-dependent cell death in several cancer cell lines and to shrink tumors *in vivo* ^11,36^. To estimate the cytoxicity of PpIX in CLL cells, we conducted a 72-hour cell viability assay using the wt-p53 EHEB B-CLL cell line and the p53-null HL60 acute promyelocytic leukaemia cells. 1 uM RITA (Reactivation of p53 and Induction of tumor cell apoptosis) was used for comparison between two compounds known to activate the p53 pathway. Cisplatin (CDDP) was used as a reference compound. The viability of EHEB and HL-60 cells was significantly reduced with the increasing concentrations of PpIX (**Figure 2A, 2B**). Of note, PpIX did not inhibit the proliferation of PBMCs at the concentrations effective in cancer cells (**Figure 2C**). Based on the above and the IC_50_ values, which are 2.5 μg/ml for EHEB and 2.4 μg/ml for HL60, we next investigated the mechanism of growth inhibition of cancer cells by PpIX and applied 2.5 μg/ml dose in all further experiments.

**Figure 2.**
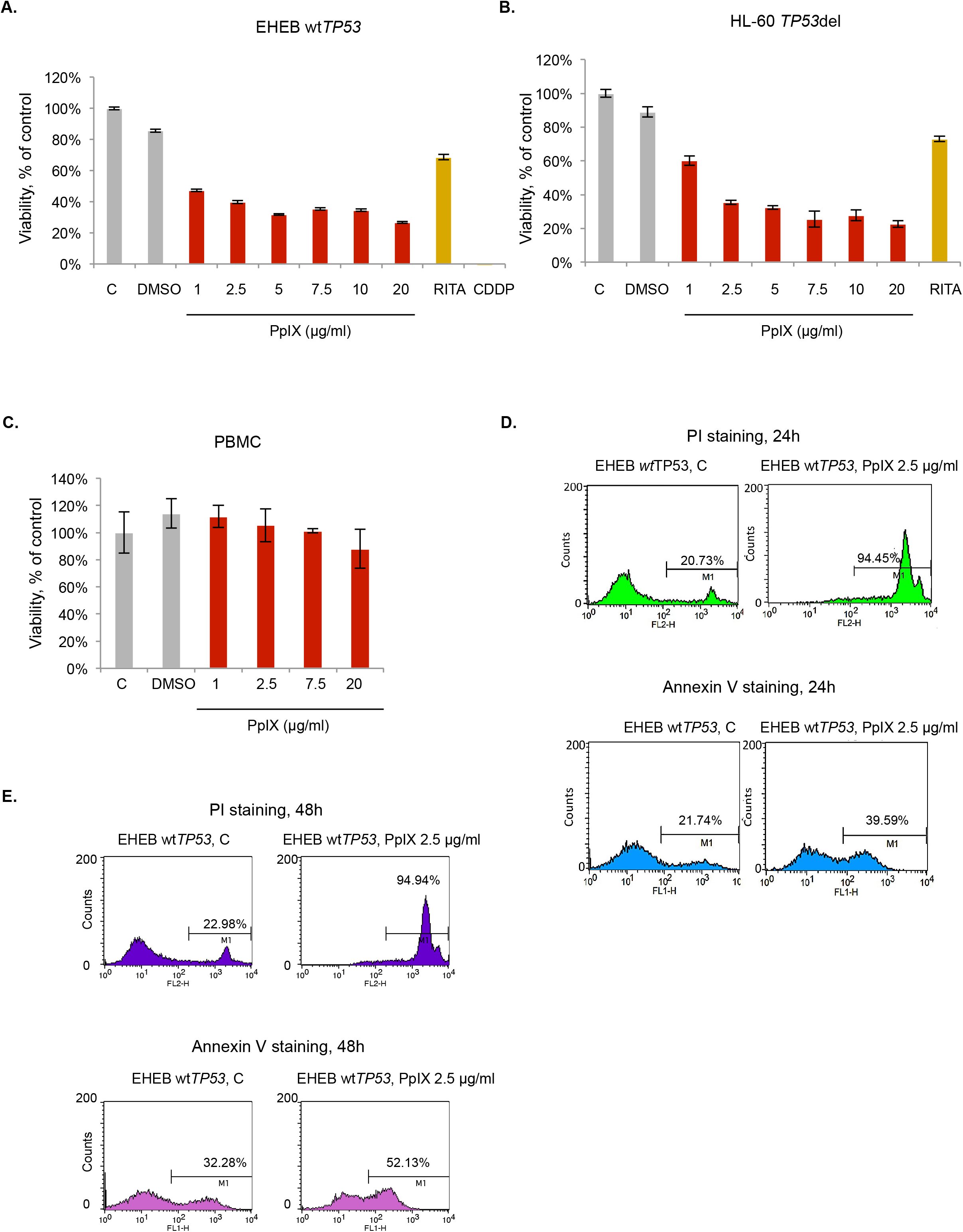
Protoporphyrin IX inhibits proliferation and induces apoptosis in chronic B cell leukaemia cells without affecting normal cells. **A.** Viability test (MTT) shows inhibition of proliferation of EHEB cells by PpIX (red bars), RITA (1 μM) (dark yellow bar) and cisplatin (50 μM) (CDDP, yellow bar) after 72h treatment. DMSO was used at 0.5%. Error bars represent SD values. n=3, C – untreated control **B.** Viability assay (MTT) in HL60 cells treated with increasing PpIX doses (red bars) or RITA (1 μM) (dark yellow bars) for 72h. DMSO was used at 0.5%. Error bars represent SD values. n=3, C – untreated control **C.** Viability of PBMCs treated with PpIX for 72 hours (red bars). DMSO was used at 0.5%. The error bars represent SD values. n=3, C – untreated control **D.** PpIX (2.5 μg/ml) activates late (upper panel) and early apoptosis (lower panel) in EHEB cells after 24h and 48h treatment **(E).** C – control treated with DMSO

To assess the induction of cancer cell death, cells were treated with PpIX and stained with propidium iodide (PI) or Annexin V. Compared with untreated control cells, PpIX-treated EHEB cells showed a significant increase in the PI positive population (late apoptosis), from 20.73% to 94.45% and in the Annexin V positive population (early apoptosis), from 21.74% to 39.59% (**Figure 2D**) after 24h. After 48h treatment with PpIX, the increase in the PI positive population was from 22.98% to 94.94% and in the Annexin V positive population from 32.28% to 52.13% (**Figure 2E**). We did not detect any increase of PBMCs in subG1 phase after treatment with 5 or 10 μg/ml PpIX, corroborating the lack of toxicity induction by PpIX in normal cells (**Supplementary Figure 1A & B**). The results in PBMCs correlated with the lack of induction of growth inhibition when treated with PpIX as assessed by MTT assay suggesting a redundant role of PpIX in healthy PBMCs (**Figure 2C**).

#### PpIX induces p53- and TAp73-dependent apoptosis in wt-p53 CLL cells but not in PBMCs

To determine the mechanism of induction of apoptosis by PpIX in CLL cells, EHEB cells were treated with 2.5 μg/ml PpIX for 8h and analysed for the expression of p53 and TAp73 apoptotic target genes *BAX* and *PUMA* by quantitative PCR (qPCR). Induction of *BAX* and *PUMA* on mRNA levels was detected (**Figure 3A**) and this corresponded to the accumulation of BAX and PUMA protein levels as assessed by western blots (**Figure 3B, C**). As reported previously, consistently with the induction of BAX and PUMA, PpIX induced p53 and TAp73 levels in EHEB cells in a time- and dose-dependent manner (**Figure 3B, C**). Interestingly, p53 accumulated more rapidly comparing to TAp73, suggesting preferential binding of PpIX to p53 in EHEB cells (**Figure 3C**). Induction of p53 preceded the accumulation of the apoptotic protein BID and cleaved PARP. In healthy PMBCs, 2.5 μg/ml PpIX induced p53 levels. However, it did not correlate with the induction of apoptotic p53 targets PUMA and BID (**Figure 3D**). In cancer cells, in addition to *PUMA* and *BAX*, PpIX induced the expression of heme oxygenase *HMOX-1* (**Figure 3A**), a stress response gene. This is in agreement with the previous studies showing that PpIX induces antioxidant response in cancer cells^12^. PpIX is an inhibitor of thioredoxin reductase (TrxR), a selenoprotein that plays a critical role in the oxidoreductase system and is often overexpressed in cancers. Inhibition of TrxR by PpIX induces reactive oxygen species (ROS) in cancer cells, triggers antioxidant response and contributes to cancer cell death^12,40^. Thus, we speculate that PpIX induces p53- and TAp73-dependent apoptosis and ROS in B-CLL cells without affecting non-transformed cells. The putative mechanism of leukaemia cells’ death is via stabilisation of p53 and TAp73 resulting from the inhibition of their interactions with oncogenic MDM2 and MDM4, the parallel accumulation of ROS resulting from inhibition of TrxR and induction of potent apoptosis (**Figure 3E**).

**Figure 3.**
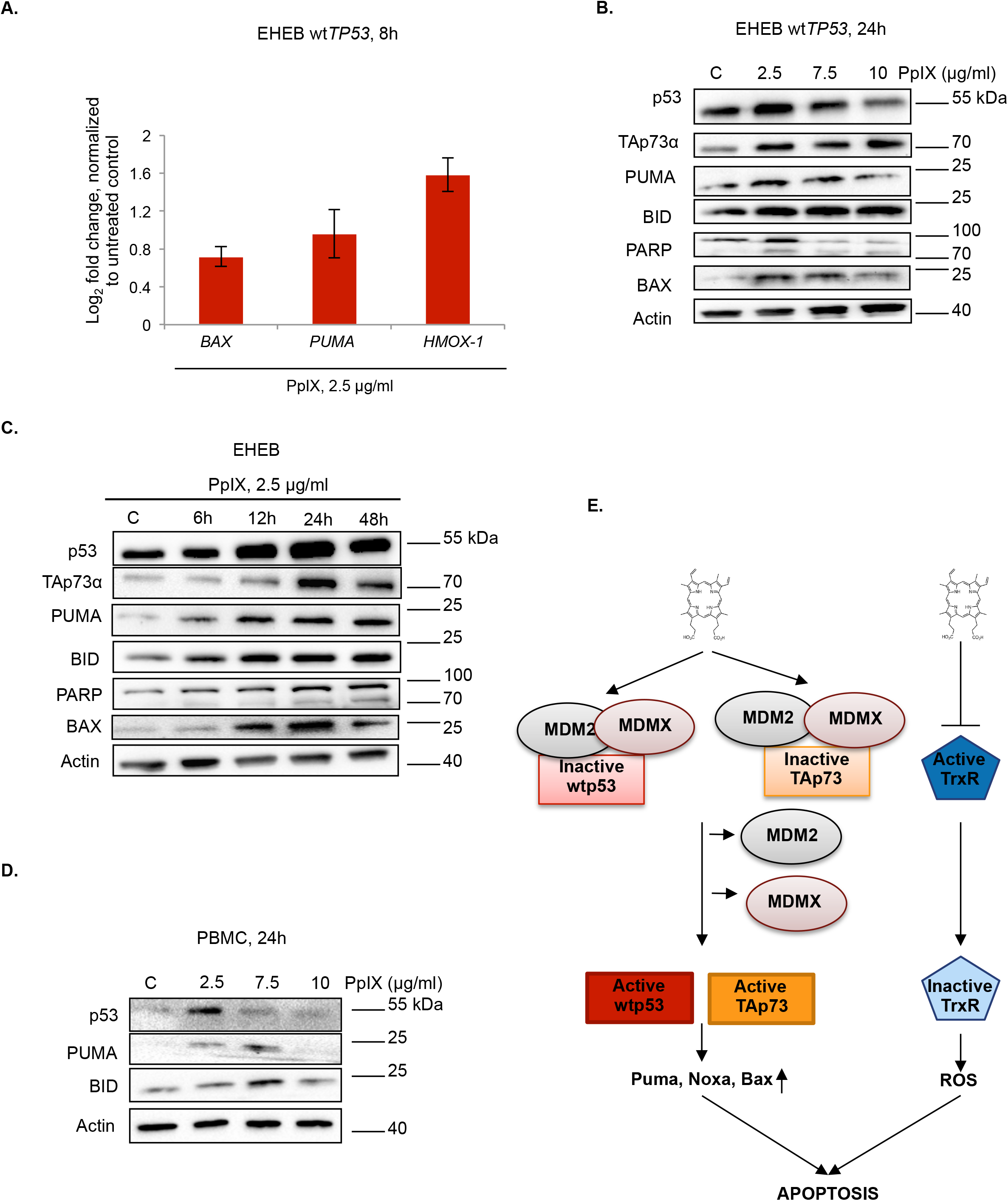
PpIX induces p53- and p73-related apoptosis in EHEB cells. **A.** PpIX (2.5 μg/ml) induces expression of apoptotic genes *BAX, PUMA* and antioxidant response gene *HMOX-1* in EHEB leukaemia cells. **B.** Western blot analysis of EHEB cells treated with increasing doses of PpIX for 24h demonstrates accumulation of p53 and TAp73 and p53-downstream apoptotic proteins. C – untreated control **C.** PpIX (2.5 μg/ml) induces p53 and TAp73 and p53-downstream apoptotic targets in EHEB cells in a time dependent manner. C – untreated control **D.** Western blot analysis of PBMCs treated with increasing doses of PpIX for 24h. C – untreated control **E.** A scheme representing the mechanism of induction of cell death in CLL cells by PpIX. PpIX simultaneously disrupts interactions between p53 or TAp73 and oncogenic MDM2 and MDM4. This results in p53 and TAp73 accumulation, induction of p53-downstream apoptotic proteins and cancer cell death.

## Discussion

Genetic or pharmacological restoration of wt-p53 activity leads to regression of tumors *in vivo* ^41–43^. Several therapeutic strategies have been developed to activate the p53 protein in tumors depending on the *TP53* gene status. Thus, in cancers, harbouring missense *TP53* gene mutations, small molecules belonging to the group of Michael acceptors that target reactive cysteine groups in the p53 core domain were found to be highly effective. These compounds refold mutant p53 protein to wild-type like conformation and induce p53-dependent apoptosis both in cancer cells *in vitro* and *in vivo*^44–47^. The most promising compound, a small molecule APR-246, a pro-drug converted to methylene quinuclidinone (MQ) is currently tested in Phase II trial in TP53-mutated high-grade serous ovarian cancer in combination with carboplatin and pegylated liposomal doxorubicin hydrochloride (PLD) (Clinical trial identifier: NCT02098343).

In cancers expressing wt-p53 protein, p53 pathway is often inhibited by the protein-protein interactions. MDM2 protein is a major E3 ubiquitin ligase of p53, which binds to p53 in the absence of cellular stress. The binding is *via* the N-terminal transactivation domain of p53 and the central domain of MDM2, inhibits p53 transcription activity and drives p53 to ubiquitin-dependent degradation. In addition, MDM2 promotes its self-degradation as well as of its homolog, MDM4 protein. MDM4, unlike MDM2, does not degrade p53 but binds to p53 N-terminus and inhibits its transcription activity. MDM2 and MDM4 are often overexpressed in cancers and therefore serve as promising therapeutic targets^20,43,48^. A large data of evidence shows that small-molecule antagonists of MDM2 restore the activity of wt-p53 in cells and thus are promising candidates for improved therapy of wtTP53 cancers.

Several compounds that bind to MDM2 and inhibit p53/MDM2 interactions are in pre-clinical and clinical development^49–51^. However, the high-affinity inhibitors of MDM2 might not be applicable in the clinical practice due to the development of resistant mutations. A recent study showed that prolonged exposure of wt-p53 harbouring cancer cells to RG7388 (idasanutlin) leads to the development of resistant mutations in the *TP53* gene^52^. Several clinical studies with a small molecule MDM2 inhibitor, AMG-232, which binds with picomolar affinity to MDM2^53^ are on-going; however, the compound is not approved yet and targets only MDM2 oncoprotein, without displaying inhibitory effect towards MDM4.

Studies have shown that PpIX, induces apoptosis in cancer cells, binds to p53 and TAp73 and induces p53-downstream apoptotic genes^11,36,37^. Cancer cells harbouring *wtTP53* gene, undergo cell death via a p53-dependent mechanism as PpIX inhibits p53/MDM2 complexes which leads to p53 stabilisation and a subsequent induction of apoptosis^36^. Next, PpIX inhibits TAp73/MDM2 and TAp73/MDM4 interactions, which induces TAp73 accumulation and TAp73-dependent apoptosis in cancers with *TP53* gene deletions or mutations^12,29^. This is in agreement with previous studies showing that colon cancer cells harbouring wtTP53 gene are more sensitive to photodynamic therapy with Photofrin®, a derivative of hematoporphyrin and, a close structural analog of PpIX^54^. Based on the above and our observations, we speculated that the pre-treatment of cancer cells with porphyrin-like photosensitizers activate the p53 pathway via p53 stabilisation which sensitizes cancer cells to photodynamic reaction induced by light matching the absorption spectrum of the compound. Here we show, using fluorescent two-hybrid technology, yeast-based p53 reporter assay and pull-down that PpIX inhibits p53/MDM2 and p53/MDM4 interactions.

Thus, PpIX, which unlike nutlin binds to p53 and not to MDM2, is a novel dual inhibitor of p53/MDM2 and p53/MDM4 interactions. It is particularly interesting that it has been demonstrated that the combined inhibition of MDM2 and MDM4 led to the enhanced p53 response and tumor regression in virus-positive Merkel cell carcinoma^55^.

Previous studies showed that PpIX induces apoptosis in human lung and pancreatic cancer cells by activating TAp73. However, the potency of the compound against haematological tissue cancers has not been tested. We applied B-CLL cell line EHEB, harbouring *wtTP53* gene and showed the induction of apoptosis by PpIX at concentrations that have no effect on normal PBMCs. The p53-null HL60 cells were also sensitive at the tested doses; however, the induction of apoptosis has not been unequivocally studied. Realtime PCR and western blot analysis revealed that PpIX induces expression of p53-downstream apoptotic *PUMA* and *BAX*. This was in agreement with simultaneous induction of p53 and TAp73 levels and accumulation of the cleaved PARP. Thus, we conclude that PpIX stabilizes both p53 and TAp73 by targeting their interaction with oncogenic MDM2 and MDM4. In addition, induction of *HMOX-1* stress response gene, suggests induction of ROS in EHEB cells. We, alongside others, showed that PpIX is an inhibitor of thioredoxin reductase, a key enzyme of the thioredoxin antioxidant defence system^12,40,56^. Inhibition of TrxR leads to induction of ROS, thus we speculate that PpIX induces *HMOX-1* in EHEB cells by activating stress response pathway due to inhibition of TrxR. Of note, several p53 targeting compounds have already been shown to inhibit TrxR which resulted in potent cancer cells eradication^57,58^. Thus, simultaneous targeting of cancer cell vulnerabilities by PpIX; namely, inactive tumor suppressors and oncogenic TrxR might bring a selective advantage over compounds already in clinical development. This is due to the fact that targeting several apoptosis-promoting pathways by PpIX might drastically reduce the risk of development of treatment resistance, as evidenced for the approved targeted therapies. Next, it has become apparent from our study that PpIX dose not induce apoptosis in healthy PBMCs. Even though, p53 was upregulated after 24h, we did not detect accumulation of apoptotic proteins PUMA and BID. This was in agreement with the lack of growth inhibition of PBMCs by PpIX as assessed by the MTT assay. Taken together, PpIX, unlike approved modalities, might have very little effect on healthy bone marrow cells, thus making the compound particularly attractive for the management of paediatric and elderly blood cancers. The best clinical outcome in the management of leukaemia is achieved in patients undergoing haematopoietic stem cell transplantation (HSCT). However, conditioning with busulfan alone or in combination with other myeloablative drugs is aggressive^59^, has a low therapeutic window and despite good protective effects of N-acetyl-l-cysteine (NAC)^60^ the side effects, particularly in older patients, result in low survival rates. In addition, busulfan has been demonstrated to affect fertility in female survivors of childhood cancers^61^. Infants and very young children undergoing haematopoietic stem cells transplantation (HSCT) are a vulnerable group of patients and are particularly sensitive to HSCT-related morbidities^62^. Next, late cardio-toxicities of anthracycline and anthraquinone frequently used in HSCT is often a significant health burden in childhood cancer survivors^63^. Thus, new treatment strategies are needed to treat paediatric leukaemia patients.

PM2, a stapled peptide that binds to MDM2 and MDM4 and stabilizes p53 has been recently described^64^ and a small molecule, LEM2, inhibiting both TAp73/MDM2 and TAp73/MDM4 interactions has been discovered in a yeast-based reporter assay^65^ However, in comparison to PpIX, the capacity of these compounds in activating both p53 and TAp73 have not been unequivocally tested.

The exact mechanism of how PpIX inhibits p53 and p73 interactions with MDM2 and MDM4 remains to be elucidated. We speculate that by binding to p53 or p73 N-terminus, PpIX might alter the conformation of the α-helix spanning the MDM2 binding residues F19, W23, and L26, which might in turn prevent the interaction between p53/MDM2 and p53/MDM4.

Summing up, our study demonstrates that PpIX is a potent activator of p53 and TAp73 in B-CLL. Our findings might speed up repurposing of PpIX in treating cancers containing wt-p53 and TAp73 and with high expression of MDM2 and MDM4 oncogenes.

## Authors contribution

LY, NM and PA, performed experimental work and drafted figures except from Figure 1a-c and LY and NM drafted the manuscript, VA set up conditions and drafted figure for yeast based assay, JZP initiated the project, supervised the study, selected the methods and conditions, obtained the funding, prepared final figures, drafted the manuscript, prepared final version and was responsible for the contact with the Journal.

All authors read and approved the final version.

## Acknowledgements

This work was supported by grants to J.Z.P. from Stockholm Läns Landsting, Karolinska Institutet, Karolinska Institutet/MD Anderson Cancer Center Sister Institution Network Fund (SINF), The Strategic Research Program in Cancer Karolinska Institutet (StratCan) and by Amgen Program (NM). We thank Sir David Lane and Dr Galina Selivanova for helpful discussions and suggestions. Special thanks are addressed to Dr Alberto Inga and Dr Virginia Andreotti. We would also like to thank Dr Alicja Sznarkowska, Dr Elisabeth Hedström and Dr Suhas Dereker for helpful feedback. The authors are grateful to Yari Ciribilli for technical assistance and to Dr Kourosh Zolghadr from ChromoTek GmbH for helpful feedback on F2H® studies. We are greatly indebted to all our colleagues who shared with us their reagents and cells.

---

**Figure S1 related to Figure 2. PpIX does not induce apoptosis in PBMCs after 16h treatment.**

**A.** Histograms showing cell cycle distribution of PI-stained PBMCs treated with PpIX for 16h. C – control treated with DMSO

**B.** Graph showing the percentage of PBMC cells accumulated is subG1 phase after PpIX treatment for 16h. n=3, error bars represent SD values.

