## Supplementary figures and images for "Protoporphyrin IX is a dual inhibitor of p53/MDM2 and p53/MDM4 interactions and induces apoptosis in B-cell chronic lymphocytic leukaemia cells"

### Supplemental Figure 1

**Supplementary Figure 1**

**A.**

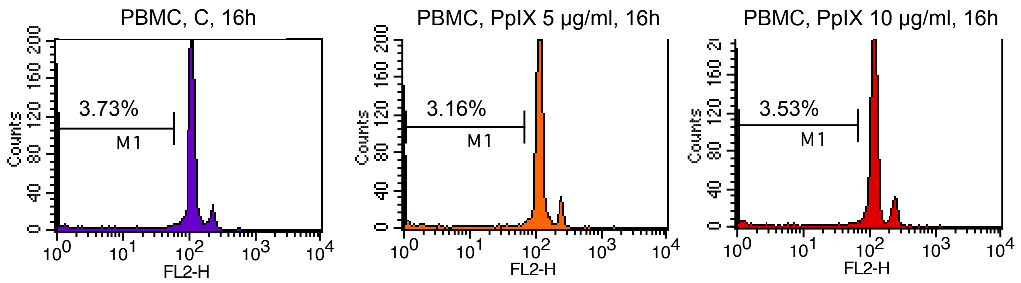


**B.**

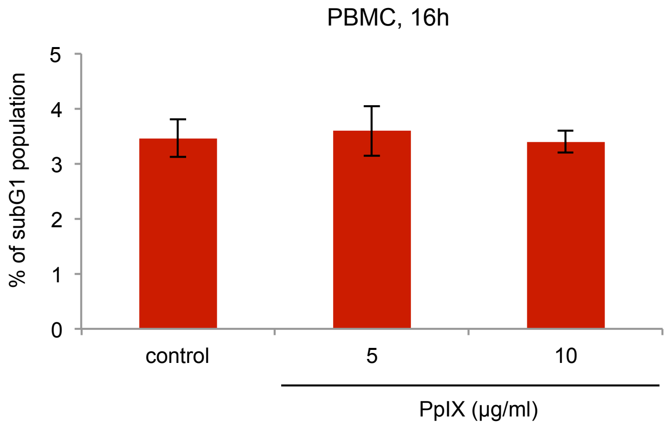
